# Mobilome-driven segregation of the resistome in biological wastewater treatment

**DOI:** 10.1101/2021.11.15.468621

**Authors:** Laura de Nies, Susheel Bhanu Busi, Benoit Josef Kunath, Patrick May, Paul Wilmes

**Affiliations:** Luxembourg Centre for Systems Biomedicine, 7, avenue des Hauts-Fourneaux, Esch-sur-Alzette, L-4362, Luxembourg; Department of Life Sciences and Medicine, Faculty of Science, Technology and Medicine, University of Luxembourg, 6, avenue du Swing, Belvaux, L-4367, Luxembourg

**Author notes:** Corresponding author: Paul Wilmes.

## Abstract

Biological wastewater treatment plants (BWWTP) are considered to be hotspots of evolution and subsequent spread of antimicrobial resistance (AMR). Mobile genetic elements (MGEs) promote the mobilization and dissemination of antimicrobial resistance genes (ARGs) and are thereby critical mediators of AMR within the BWWTP microbial community. At present, it is unclear whether specific AMR categories are differentially disseminated via bacteriophages (phages) or plasmids. To understand the segregation of AMR in relation to MGEs, we analyzed meta-omic (metagenomic, metatranscriptomic and metaproteomic) data systematically collected over 1.5 years from a BWWTP. Our results showed a core group of fifteen AMR categories which were found across all timepoints. Some of these AMR categories were disseminated exclusively (bacitracin) or primarily (aminoglycoside, MLS, sulfonamide) via plasmids or phages (fosfomycin and peptide), whereas others were disseminated equally by both MGEs. Subsequent expression- and protein-level analyses further demonstrated that aminoglycoside, bacitracin and sulfonamide resistance genes were expressed more by plasmids, in contrast to fosfomycin and peptide AMR expression by phages, thereby validating our genomic findings. Longitudinal assessment further underlined these findings whereby the log2-fold changes of aminoglycoside, bacitracin and sulfonamide resistance genes were increased in plasmids, while fosfomycin and peptide resistance showed similar trends in phages. In the analyzed communities, the dominant taxon *Candidatus* Microthrix parvicella was a major contributor to several AMR categories whereby its plasmids primarily mediated aminoglycoside resistance. Importantly, we also found AMR associated with ESKAPEE pathogens within the BWWTP, for which MGEs also contributed differentially to the dissemination of ARGs. Collectively our findings pave the way towards understanding the segmentation of AMR within MGEs, thereby shedding new light on resistome populations and their mediators, essential elements that are of immediate relevance to human health.

## Introduction

Throughout human history, bacterial infections have been a major cause of both disease and mortality^1^. The discovery as well as the subsequent development and medical use of antibiotics have provided effective treatment options which limited the development and spread of bacterial pathogens. However, the use of antibiotics has exacerbated the emergence of antimicrobial resistance (AMR) in both commensal and pathogenic bacteria^2^. As a result, AMR, as the “silent pandemic”, has become a prevalent threat to human health^3–5^.

From a public health perspective, biological wastewater treatment plants (BWWTPs) are considered hotspots of AMR due to the convergence of antibiotics with resistant, potentially pathogenic microorganisms originating from both the general population as well as agriculture and healthcare services^6,7^. Additionally, the mobilization of antimicrobial resistance genes (ARGs) through rampant horizontal gene transfer (HGT) promotes the dissemination of AMR within the BWWTP microbial community^8^. Therefore, BWWTPs represent an environment exceptionally suited for the evolution and subsequent spread of AMR^9,10^. To date, more than 32 studies have documented the role of BWWTPs as key reservoirs of AMR^11^. Furthermore, BWWTPs generally do not contain the necessary infrastructure to remove either ARGs or resistant bacteria, which are released into the receiving water via the effluent, promoting its spread in the environment at large^12^. Most often these are surface water bodies such as rivers, which contribute to the further dissemination of AMR and resistant bacteria among environmental microorganisms^13^. Acquired resistance may in turn be carried over to humans and animals using these water resources. In fact, there is strong evidence suggesting that ARGs from environmental bacteria can be taken up by human-associated and pathogenic bacteria^14,15^. From an epidemiological and surveillance perspective, BWWTPs also provide samples representative of entire populations^16^. As such, BWWTPs have recently been crucial for the monitoring of SARS-CoV-2 within the human population^17^. Overall, to increase our understanding of the dissemination of AMR and the underlying mechanisms as well as its general prevalence, it is necessary to map the resistome of various environments starting with biological BWWTPs because it is critical to unravel the extent to which they act as reservoirs for the dissemination of antimicrobial resistance genes (ARGs) to bacterial pathogens. Moreover, understanding the community-level overviews of the ARG potential and its expression, coupled with population-level linking, including to pathogens, may allow for efficient monitoring of pathogenic and AMR potential with broad impacts on human health.

The conditions such as the presence of resistance genes^18^, and sub-inhibitory antibiotic selection pressure from various sources^8^ facilitate HGT of ARGs into new hosts through the mobilome, i.e. mobile genetic elements (MGEs). Acquisition of ARGs via MGEs primarily occurs through two mechanisms: conjugation or transduction^19^. In conjugation, plasmids carrying one or more resistance genes are transferred between microorganisms^20^, while in transduction bacteriophages carrying ARGs infect bacteria and integrate their genome into those of the host thereby conferring resistance^21^. Of these mechanisms, conjugation is often thought to have the greatest influence on the dissemination of ARGs, while transduction is deemed less important^8^. In general terms, studies concerning AMR and its dissemination focus either on phage^22,23^ or plasmids solely^24^. Alternatively, the two are treated collectively^12,25^ without a comprehensive comparative analysis. This circumstance has created a knowledge gap whereby the contributions of plasmids and phages as independent entities to AMR transmission within complex communities, such as those found in biological BWWTPs, is largely unknown.

To shed light on the evolution, dissemination and potential segregation of AMR within MGEs in a WTTP microbial community, we leveraged longitudinal meta-omics data (metagenomics, metatranscriptomics and metaproteomics). Samples collected for 51 consecutive weeks over a period of 1.5 years, were used to characterize the resistome. We found that several bacterial orders such as Acidimicrobiales, Burkholderiales and Pseudomonadales were associated with 29 AMR categories across all timepoints. Our longitudinal analysis demonstrated that MGEs are important drivers of AMR dissemination within BWWTPs and that assessing the activity of the ARGs is critical for understanding the underlying mechanisms. More importantly, we reveal that MGEs, i.e. plasmidomes and phageomes, contribute differentially to AMR dissemination. Furthermore, we observed this phenomenon in clinically-relevant taxa such as the ESKAPEE pathogens^26^, for which plasmids and phages were exclusively associated with specific ARGs. Collectively, our data suggest that BWWTPs are critical reservoirs of AMR which show clear evidence for the segregation of distinct AMR genes within MGEs especially in complex microbial communities. In general, we believe that these findings may provide crucial insights into the segregation of the resistome via the mobilome in any and all reservoirs of AMR, including but not limited to animals, humans, and other environmental systems.

## Results

### Longitudinal assessment of the resistome within a BWWTP

To characterize the BWWTP resistome, we sampled a municipal BWWTP on a weekly basis over a 1.5 year period (ranging from 21-03-2011 to 03-05-2012)^27,28^. Utilizing the PathoFact pipeline we resolved the BWWTP resistome. This analysis revealed the presence of 29 different categories of AMR within the BWWTP. Subsequent longitudinal analyses highlighted enrichments in aminoglycoside, beta-lactam and multidrug resistance genes (Fig. 1a). Concomitantly, we observed specific shifts in the AMR profiles over time. For example, a shift at two timepoints (13-05-2011, 08-02-2012) highlighted a steep increase in resistance genes corresponding to glycopeptide resistance. Other AMR categories, such as diaminopyrimidine resistance, exhibited a less drastic but more fluid change in longitudinal abundance observable over multiple timepoints.

**Figure 1.**
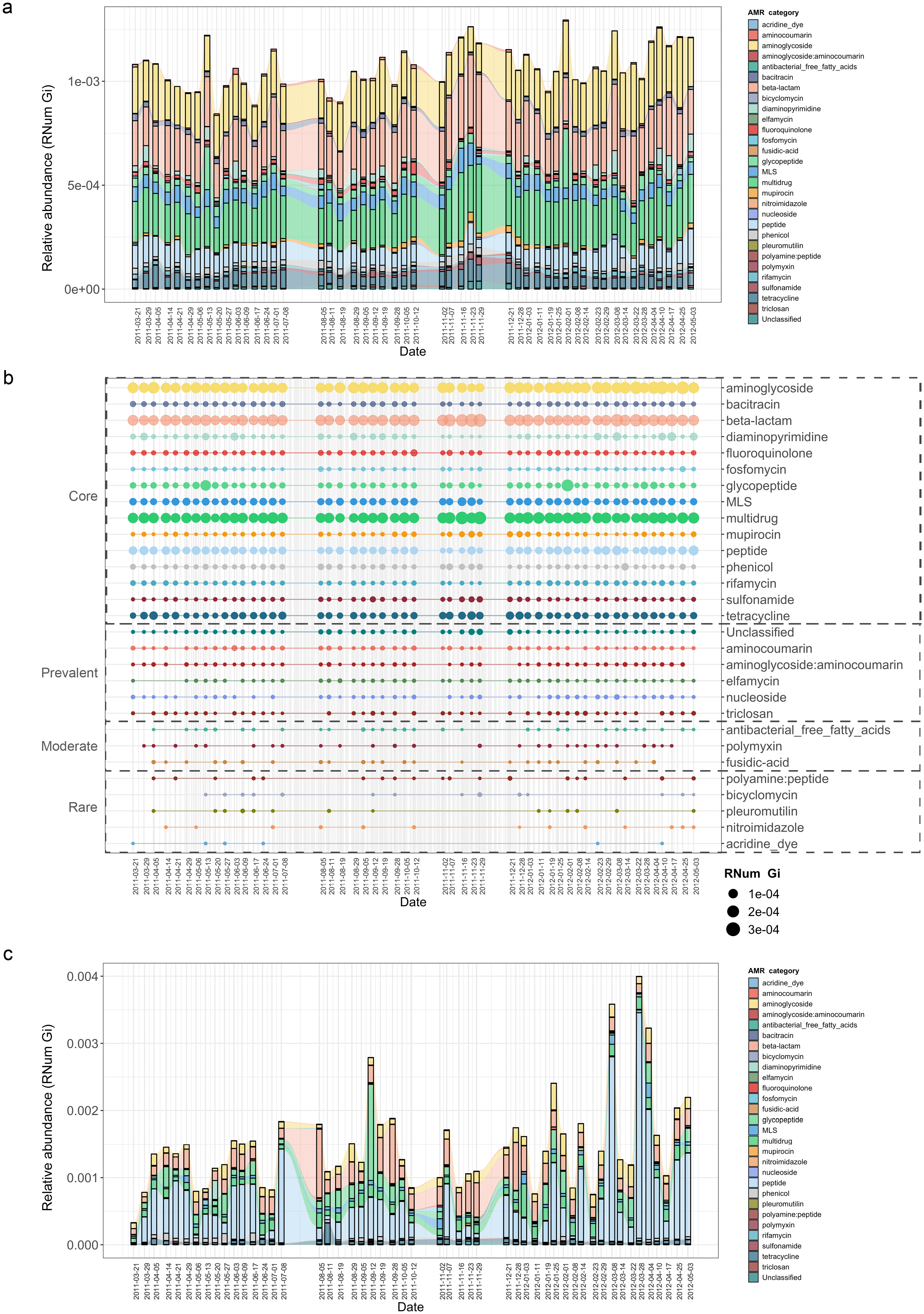
Longitudinal metagenomic and metatranscriptomic assessment of AMR a) ARG relative abundances over time within the BWWTP. b) ARG categories at various timepoints categorized in 4 distinct groups based on presence/absence: Core (all timepoints), Prevalent (>75% of timepoints), Moderate (50-75% of timepoints) and Rare (< 50% of all timepoints). c) Relative abundance levels of expressed AMR categories over time within the BWWTP.

Additionally, AMR categories were found to persist variable over time (Fig. 1b). A core group of 15 AMR categories in total were identified and found to be present across the 1.5 year sampling period. These included aminoglycoside, beta-lactam and multidrug resistance genes, which contributed the most to the pool of ARGs. A further six (aminocoumarin, aminoglycoside:aminocoumarin, elfamycin, nucleoside, triclosan and unclassified) AMR categories were found to be prevalent (>75% of all timepoints), while another three AMR categories were moderately (50 - 75% of all timepoints) present over time (Fig. 1b). Five other categories were rarely present within the BWWTP, with resistance corresponding to acridine dye only present at six of the timepoints. Altogether, this emphasized that the BWWTP resistome varies over time, substantiating the requirement for a longitudinal analysis to obtain an accurate overview of the community’s overall resistome.

Although the data thus far provided a clear overview of the BWWTP from a metagenomic perspective, it did not provide any information regarding AMR expression. We therefore utilized the corresponding metatranscriptomic dataset to investigate the expression of identified ARGs and monitor their changes, within the BWWTP, over time. In contrast to the metagenomic data, we observed a difference in AMR expression levels for several categories. Aminoglycoside, beta-lactam, and multidrug resistance identified at high levels in metagenomic information were also highly expressed within the BWWTP (Fig.1c). However, peptide resistance demonstrated the highest expression levels of all the AMR categories. We further investigated which ARG subtypes contributed to the identified peptide resistance category and found that ~90% of the expressed peptide resistance was directly contributed by a single resistance gene, *YojI* (Supp. Fig. 1), typically associated with resistance to microcins^29^ a potential adaptive strategy amongst the microbial populations in the BWWTP against these specific stressors.

### Microbial community and co-occurrence patterns of AMR

Based on the previously identified microbial community^28^, we hypothesized that the abundant and prevalent bacterial orders such as Acidimicrobiales were major contributors to the abundance in ARGs observable via metagenomics. To further investigate the contribution to AMR by the distinct microbial populations, we linked AMR genes to the contig-based taxonomic annotations of the assemblies. Herein, we identified a wide variety of taxonomic orders contributing to AMR, with multiple orders often contributing to the same resistance categories (Supp. Fig. 2). Overall, taxa belonging to Acidimicrobiales, followed by Burkholderiales, were found to encode most of the ARGs (Fig. 2a). Additionally, the abundance of ARGs linked to taxonomy varied over time. This was most noticeable during a five-week period (autumn: 02-11-2011 to 29-11-2011), where a decrease in abundance in ARGs linked to Acidimicrobiales and Bacteroidales was observed coinciding with an increase in ARG abundance in Pseudomonadales and Lactobacillales.

**Figure 2.**
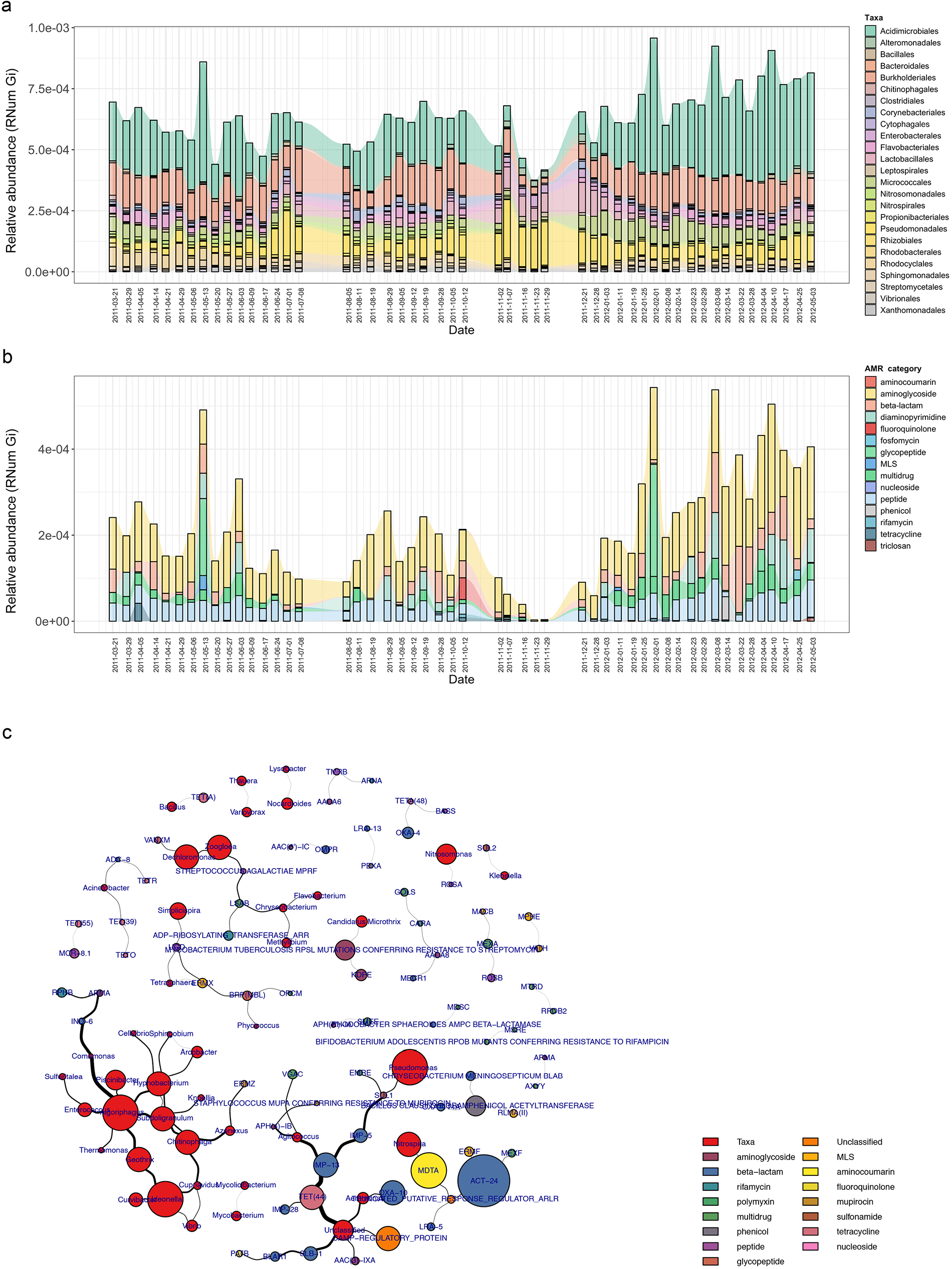
Microbial population-linked AMR a) Longitudinal ARG relative abundance levels linked to their corresponding microbial taxa (order level). b) Relative abundance of ARG categories linked to *Candidatus* Microthrix parvicella. c) Bi-partite network depicting co-occurrence patterns of individual antimicrobial resistance genes (ARGs) and microbial taxa on genus level.

Since the family Acidimicrobiales was found to be linked to the highest abundance in ARGs, we further resolved the taxonomic affiliation and identified the species *Candidatus* Microthrix parvicella (hereafter known as *M. parvicella*) to be the main contributor to AMR. *M. parvicella* was previously found to dominate this microbial community^27^ and is a well-characterized bacterium commonly occurring in the BWWTP^30^. Overall, aminoglycoside, beta-lactam, multidrug and peptide resistance were found to be abundant in this species (Fig. 2b), with aminoglycoside resistance demonstrating the highest expression levels as confirmed through metatranscriptomic analysis (Fig. 2c). Although it was not surprising to find a high abundance of ARGs linked to this species, the longitudinal variation in the abundances of these ARGs was nevertheless surprising (Fig. 2b). Furthermore, coupled to a decrease in the abundance of *M. parvicella* itself^27^, we observed an almost complete decrease in ARGs at two timepoints (23-11-2011 and 29-22-2011). However, the *M. parvicella* population recovered to levels resembling the earlier timepoints in conjunction with the abundances in ARGs towards the end of the sampling period (Fig. 2a, Fig. 2b), underlining their overall contribution to AMR within this BWWTP. Alternatively, it is plausible that the dominance of *M. parvicella* is attributable to the encoded ARGs, which in turn, may confer a fitness advantage.

In order to determine whether the abundances in ARGs may be directly associated with the community composition and population sizes over time, co-occurrence patterns between ARG subtypes and taxa (genus level) were explored using the metagenomic data. Bipartite network analyses (Fig. 2c) demonstrated that ARGs, within or across ARG types and microbial taxa, showed clear and distinct co-occurrence patterns within the BWWTP. These patterns indicated a strong segregation of distinct, taxa-specific ARG subtypes within the BWWTP community over time. One clear example was that of *M. parvicella* which encoded different aminoglycoside resistance genes (Fig. 3a). Thus, the abundance of this bacterium along with the aminoglycoside ARGs were highly correlated.

**Figure 3.**
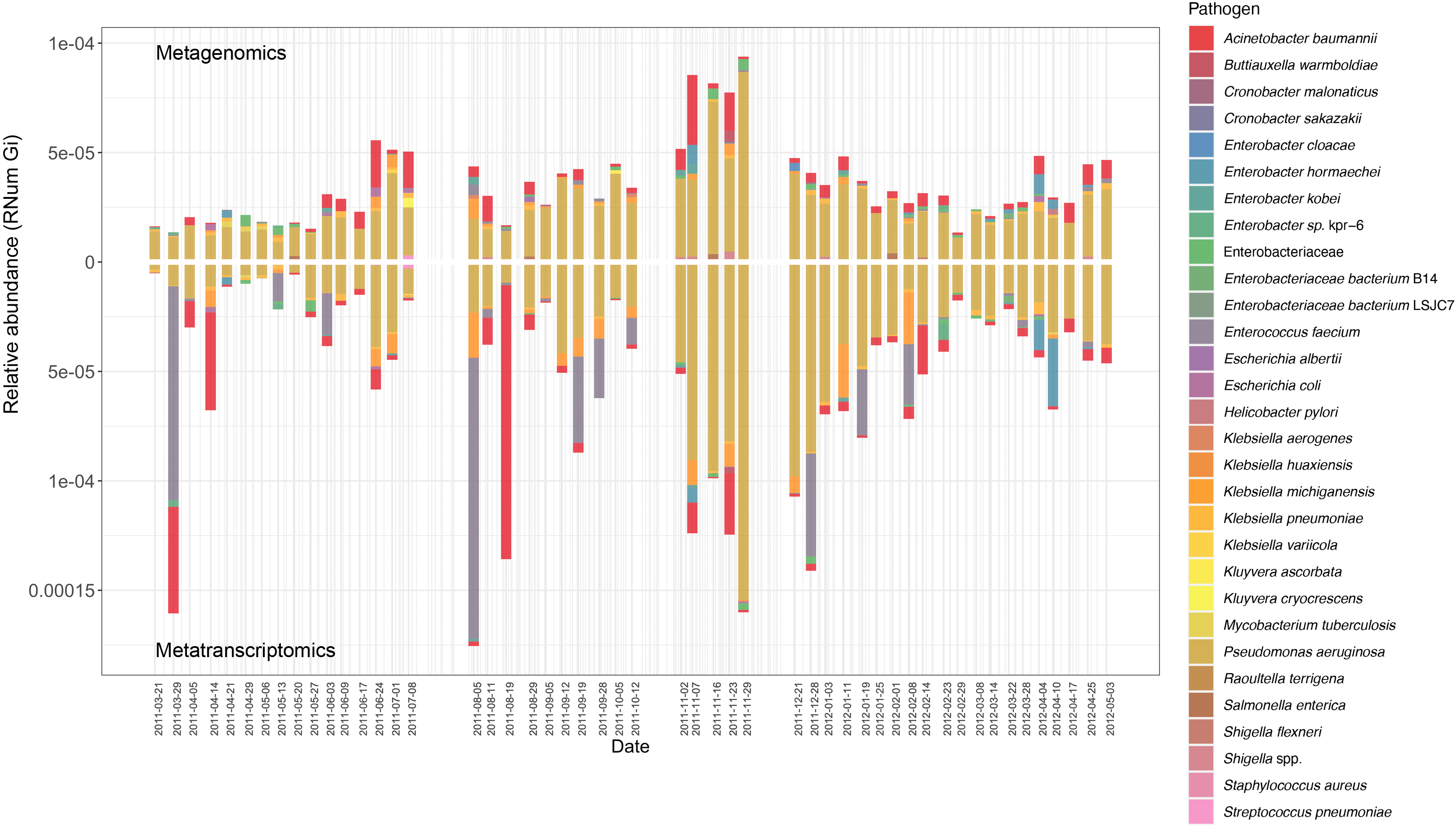
Assessment of AMR associated with clinical pathogens ARG relative abundance encoded and expressed by clinical pathogens over time within the BWWTP.

### Monitoring pathogenic microorganisms within BWWTPs

In conjunction with the families observed within BWWTPs, we also found that certain ESKAPEE pathogens^26^, such as *Klebsiella* spp. and *Pseudomonas* spp., demonstrated co-occurring patterns with ARGs (Fig. 2c).

As previously mentioned, BWWTPs represent a collection of potentially pathogenic microorganisms originating from, among others, the human population. Moreover, evidence suggests that ARGs from environmental and commensal bacteria can spread to pathogenic bacteria through HGT^19^. Therefore, we assessed the acquisition and dissemination of AMR in the extended priority list of pathogens (Table 1), characterized as such by the WHO^31^, using both metagenomics and metatranscriptomics.

**Table 1:**
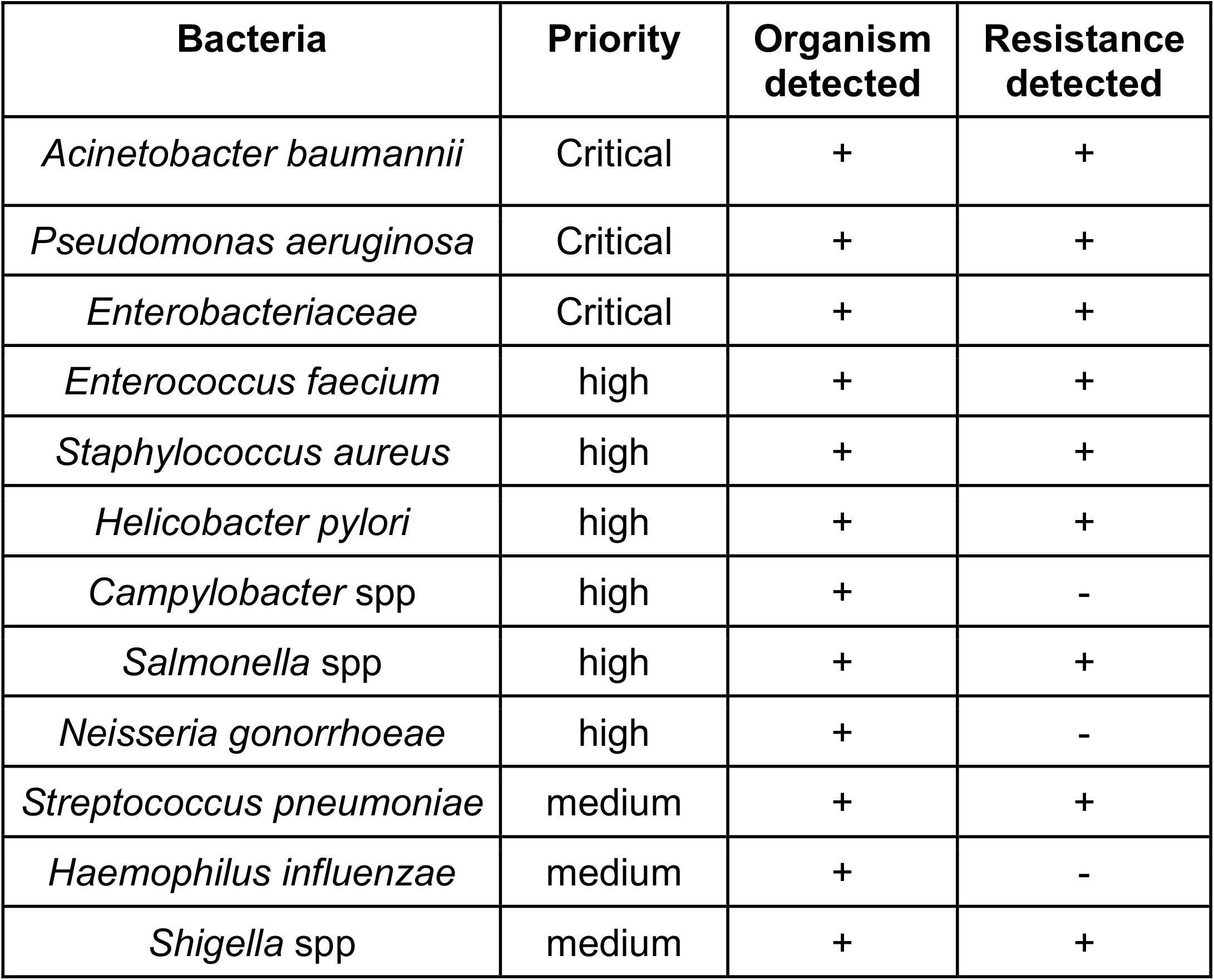
WHO priority list for research and development of new antibiotics for antibiotics-resistant bacteria^31^.

| Bacteria | Priority | Organism detected | Resistance detected |
| --- | --- | --- | --- |
| *Acinetobacter baumannii* | Critical | + | + |
| *Pseudomonas aeruginosa* | Critical | + | + |
| *Enterobacteriaceae* | Critical | + | + |
| *Enterococcus faecium* | high | + | + |
| *Staphylococcus aureus* | high | + | + |
| *Helicobacter pylori* | high | + | + |
| *Campylobacter* spp | high | + | - |
| *Salmonella* spp | high | + | + |
| *Neisseria gonorrhoeae* | high | + | - |
| *Streptococcus pneumoniae* | medium | + | + |
| *Haemophilus influenzae* | medium | + | - |
| *Shigella* spp | medium | + | + |

Of the identified pathogens (Table 1), we found that *Pseudomonas aeruginosa*, both encoded and expressed the highest abundance of ARGs, followed by *Acinetobacter baumannii*, over time within the BWWTP (Fig. 3). Moreover, an increase in ARG abundance and expression was observed in *Pseudomonas aeruginosa* during the time period, during which the otherwise dominant *M. parvicella* demonstrated reduced abundance (Fig. 2b & Fig. 3).

### Differential transmission of antimicrobial resistance via mobile genetic elements

As previously described^32,33^, the mobilome is a major contributor to the dissemination of AMR within a microbial community. Consequently, to understand (i) the role of MGE-mediated AMR transfer within the BWWTP, and (ii) to identify differential contribution of the mobilome to the dissemination of AMR, we identified both plasmids and phages within the metagenome and linked these to the respective ARGs. Overall, we found that plasmids contributed to an average of 10.8% of all ARGs, while phage contributed to an average of 6.8% of all resistance genes, confirming the general hypothesis that conjugation has the greatest influence on the dissemination of ARGs^19^. This phenomenon, however, varied across time within the BWWTP (Fig. 4a).

**Figure 4.**
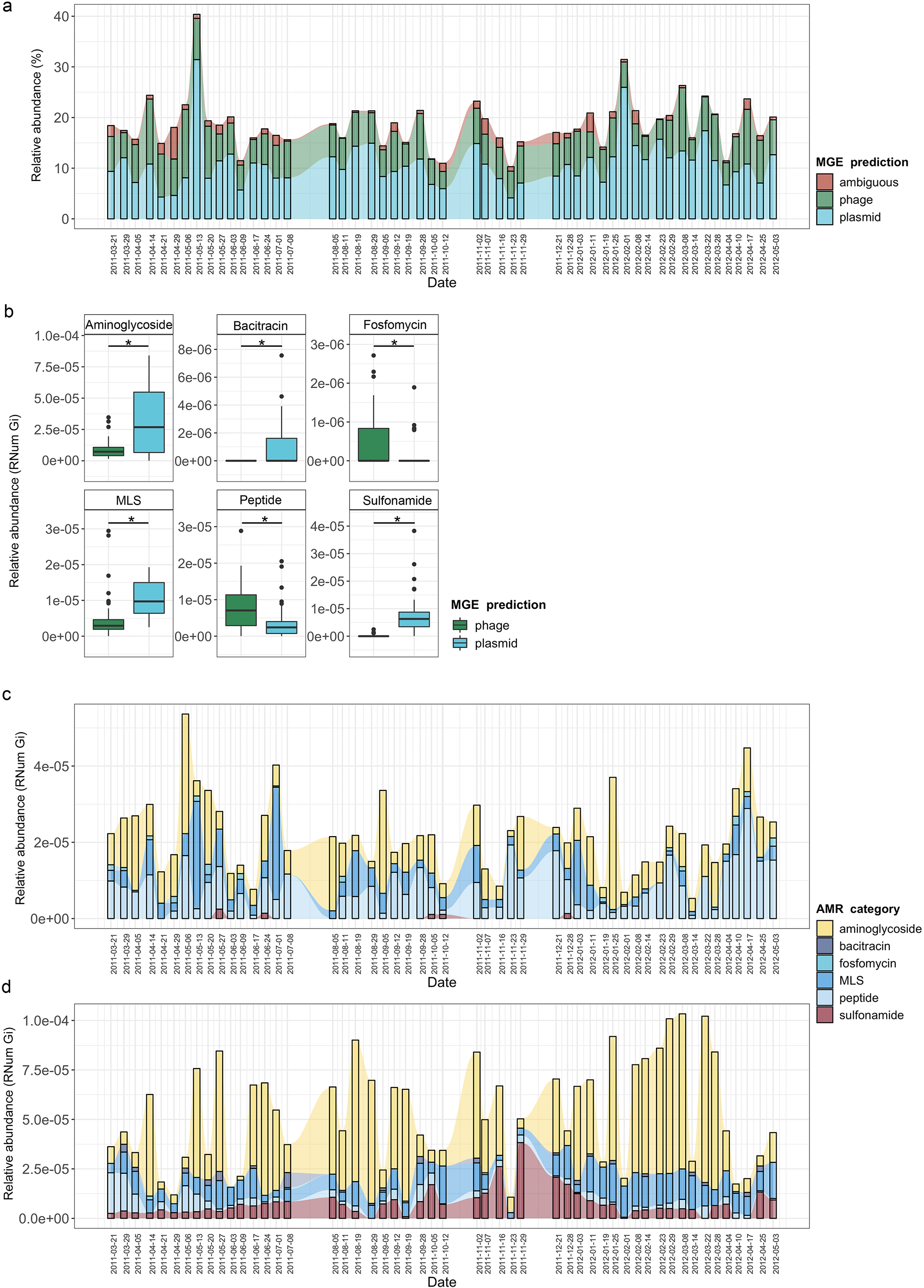
MGE-derived AMR within the BWWTP resistome a) Overall relative abundance of MGEs encoding ARGs. b) Boxplots depicting significant (*adj.p < 0.05,* Two-way ANOVA) differential abundances of ARGs encoded by plasmids vs phages. c) Relative abundance of the 6 significantly different AMR categories encoded on phages over time. d) Relative abundance of the 6 significantly different AMR categories encoded on plasmids over time.

When investigating the dissemination of AMR via MGEs, most reports typically focus on either phages or plasmids individually, or both as collective contributors to transmission^34^. To date and to our knowledge, the respective contributions of phage and plasmid to AMR transmission have not been subjected to a comprehensive comparative analysis. To facilitate a systematic, comparative view of MGE-mediated AMR, we assessed the segregation of MGEs with respect to AMR and found that phages and plasmids contributed differentially to AMR (Supp. Fig. 3). Specifically, we found a significant difference in six AMR categories when comparing ARGs encoded by phages and plasmids (Fig. 4b). Aminoglycoside, bacitracin, MLS (i.e. macrolide, lincosamide and streptogramin) and sulfonamide resistance were found to be primarily encoded by plasmids, whereas fosfomycin and peptide resistance were found to be associated with phages.

To further understand AMR in relation to the community dynamics, we investigated the abundance and segregation of the above-mentioned significant resistance categories at different timepoints within the BWWTP. We observed ARG abundances varied over time both in phages (Fig. 4c) as well as plasmids (Fig. 4d). For instance, the abundance in aminoglycoside and sulfonamide resistance, which was encoded primarily by plasmids (Supp. Fig. 4a), fluctuated widely over time in both phages and plasmids (Fig. 4c). Additionally, plasmid-mediated sulfonamide resistance was reduced at 23-11-2011, followed by its highest abundance a week later (20-11-2011), while subsequently again decreasing. Similarly, in line with the above observations, fosfomycin and peptide resistance genes, while segregating within phages, demonstrated significant fluctuations over time (Fig. 4d). In addition to the metagenome, we also contextualized the localization of the expressed ARGs within MGEs based on the metatranscriptomic information. Specifically, we found that plasmids demonstrated a significantly increased expression of aminoglycoside along with bacitracin and sulfonamide resistance genes, while the expression of glycopeptide, mupirocin and peptide resistance genes were primarily enriched in phages (Fig. 5a). These observations pertaining to plasmid-mediated AMR were in line with the metagenomic findings (Fig. 4b). Only peptide resistance was observed to be expressed via phages in contrast to the differential enrichment of fosfomycin resistance observable in the metagenomic data.

**Figure 5.**
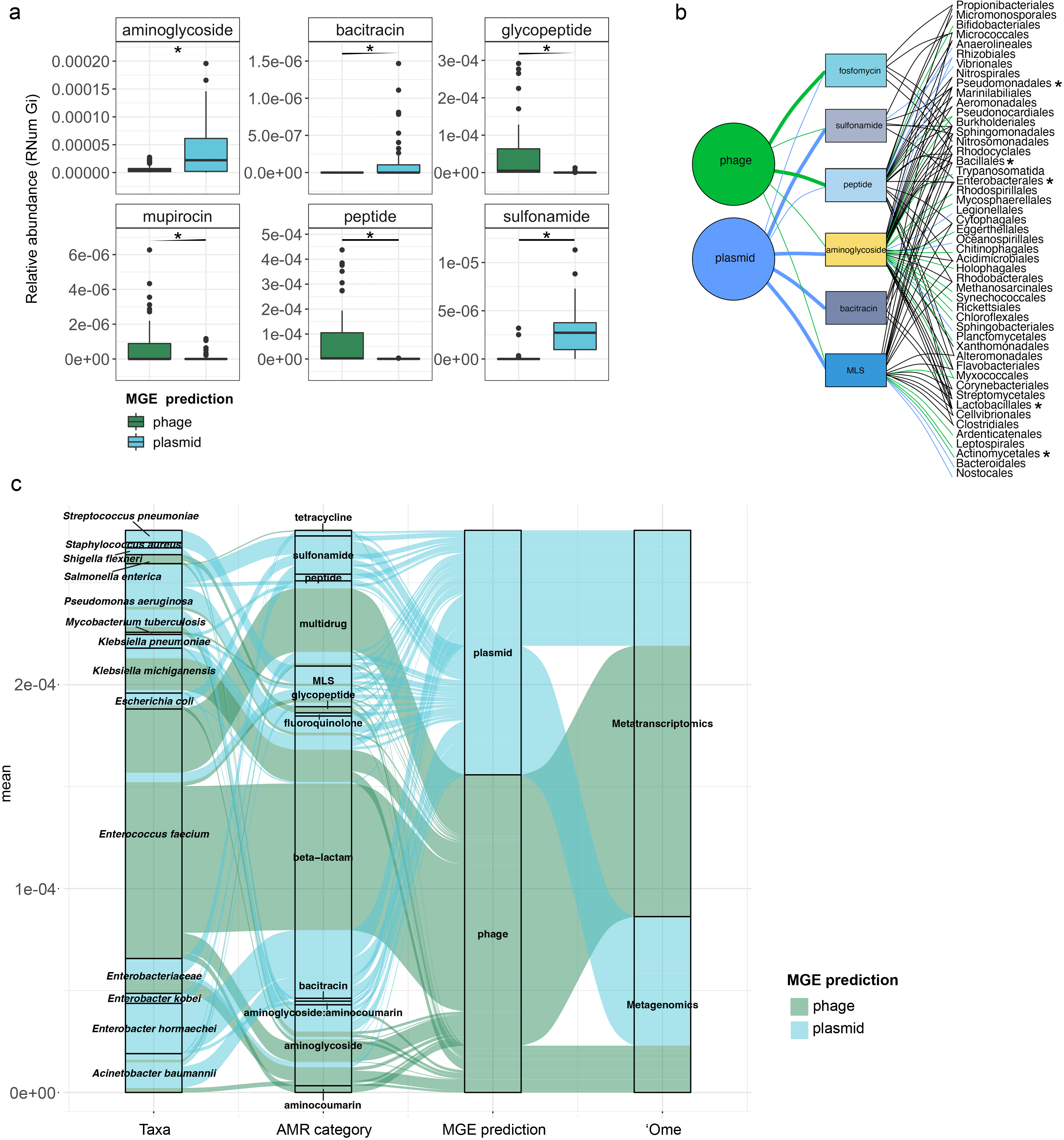
Taxonomic affiliations of MGE-derived resistance genes a) Boxplot depicting significant differential abundance (adj.p < 0.05, Two-way ANOVA) of ARGs expressed in plasmids vs phages. b) Tripartite network assessing the association of MGE-derived ARGs with the microbial taxa. Thickness of the lines representing potential niche-partitioning of the AMR category to one MGE over the other. Color of the line representing which MGE the AMR is linked to: green (phage), blue (plasmid) or black (both phage and plasmid). Asterisk denominates taxonomic orders which include known clinical pathogens. c) Alluvial plot depicting relative abundances of MGE-derived ARGs encoded (metagenome) and/or expressed (metagenome) by clinical pathogens.

### Taxonomic affiliations of MGE-derived resistance genes

When assessing the differential contributions of MGEs to AMR, we found congruency between plasmids and phages to the AMR categories and taxonomic affiliations (Fig. 5b). For example, in the metagenomic data MGEs (phage and plasmid) were predominantly associated with the same AMR category and subsequently the same taxa. However, some exceptions were observed with specific taxa associated with AMR either through plasmids or phages. For instance, MLS resistance in Bacteroidales and Nostocales was mediated solely through plasmids, whereas the same resistance category was mediated by phage in Bifidobacteriales, indicating a mechanistic basis for the segregation of AMR between taxa and MGEs.

As most bacteria harbor MGEs, we queried whether the MGE-mediated AMR categories were linked to the abundance of some of the earlier reported taxa. Interestingly, we found that peptide resistance encoded by *M. parvicella* was solely associated with phages, while aminoglycoside resistance was primarily correlated with plasmids (Supp. Fig. 4b). Other highly abundant taxa such as *Pseudomonas* and *Comamonas* (Supp. Fig. 4c-d), on the other hand, were correlated with sulfonamide resistance in addition to aminoglycoside resistance encoded on plasmids (Fig. 5b). This was further reflected within the metatranscriptome data where in taxa such as Acidimicrobiales the expression levels of aminoglycoside resistance were solely associated with plasmids (Supp. Fig. 5a). Additionally, in the Burkholderiales family, peptide and glycopeptide resistance were found to be expressed through phages (Supp. Fig. 5b).

We also found a clear segregation of the mobilome with respect to individual pathogens in the metagenome. Interestingly, plasmids were exclusively associated with AMR in six out of the fourteen relevant taxa (Fig. 5c). These included *Streptococcus pneumoniae, Staphylococcus aureus, Shigella flexneri, Klebsiella pneumoniae, Enterobacter kobei* and *Enterobacter hormaechei*. Furthermore, the plasmids were also associated with conferring peptide, multidrug, MLS, beta-lactam, fluoroquinolone, bacitracin, aminoglycoside, aminoglycoside:aminocumarin and sulfonamide resistance. Phages were exclusively associated with glycopeptide and aminoglycoside resistance *in Salmonella enterica*. Overall, our results revealed for the first time the key segregation patterns of AMR via the mobilome in taxa that are of relevance to human health and disease. Moreover, substantiating the metagenomic data, the pathogenic bacteria *S. pneumoniae, S. aureus, K. pneumoniae, E. kobei* and *E. hormaeche* were found to express ARGs solely associated with plasmids (Fig. 5c). Collectively, these findings represent an imminent threat to global health due to their potential for dissemination across reservoirs.

### Metaproteomic validation of AMR abundance and expression

In order to validate our findings with the expression (metatranscriptomic) analyses on the BWWTP, we further used the corresponding metaproteomic data to offer complementary information at the protein level. Similar to the metagenome data we found protein expression linked to aminoglycoside, beta-lactam and multidrug resistance, over time within the BWWTP (Supp. Fig. 6). Proteins linked to multidrug resistance especially were found to increase over time.

To further improve upon the understanding of the AMR expression and assess its stability across the time, we estimated the normalized protein index (NPI) per gene, as discussed in the Methods, by integrating all of the multi-omic data. The estimated NPI demonstrated stable levels of aminoglycoside and multidrug resistance within the BWWTP (Fig. 6a). Specifically, proteins conferring multidrug resistance were found to increase over time, which is in line with the gene- and expression-level observations. Furthermore, we contextualized the normalized proteins conferring AMR to their localization on MGEs. We identified five resistance categories, i.e. aminoglycoside, beta-lactam, sulfonamide, multidrug and tetracycline resistance, to be expressed through MGEs (Fig 6b). Of these categories we found that aminoglycoside resistance, in concordance with the gene and expression levels, was significantly higher mediated through plasmids compared to phages. We further found that the MGE-mediated AMR categories were associated with specific microbial taxa. with plasmid-mediated aminoglycoside resistance found to be strongly associated with the previously mentioned *M. parvicella* (Fig. 6b). On the other hand, we did not identify any peptides associated with the ESKAPEE pathogens via metaproteomics.

**Figure 6.**
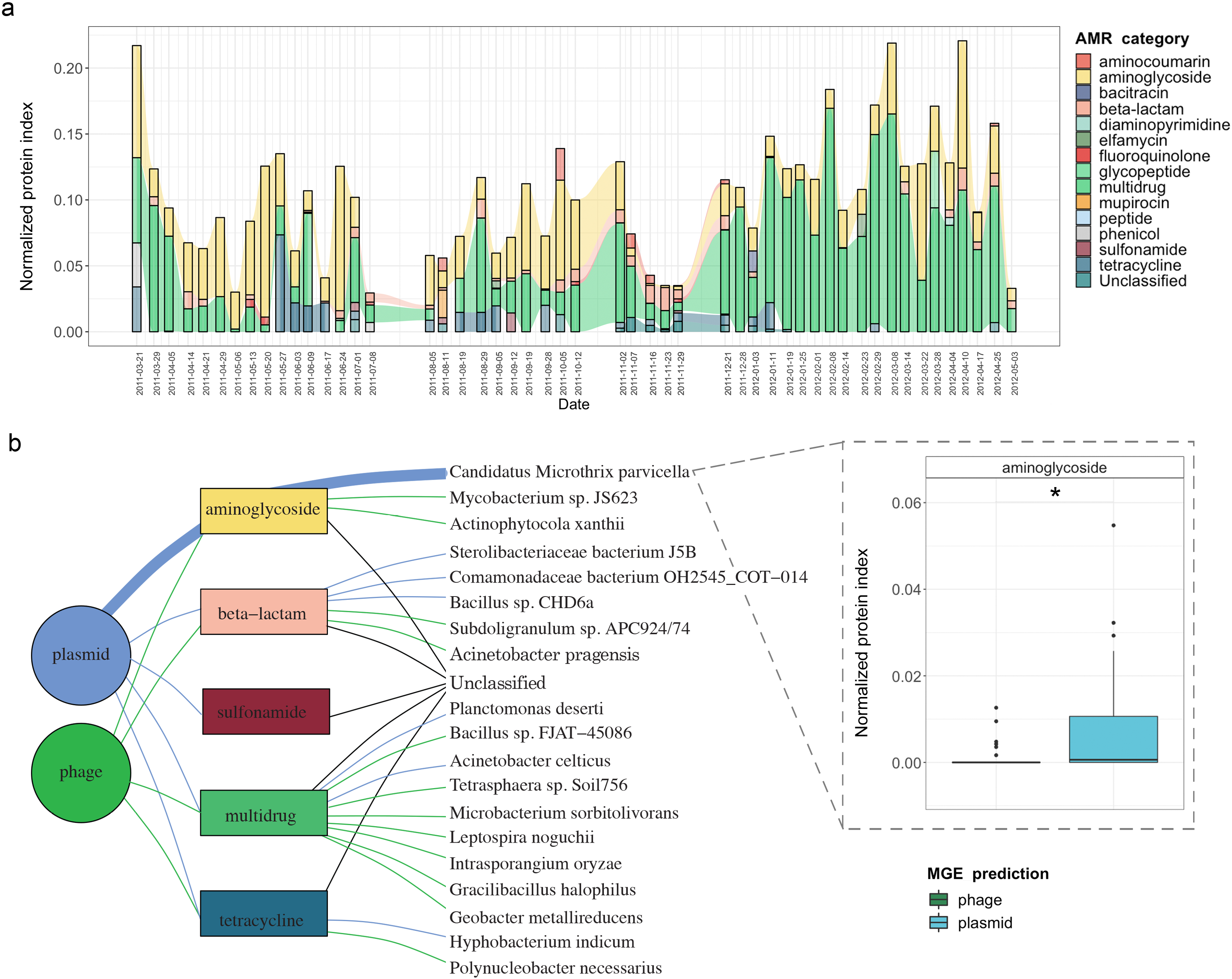
Integrative multi-omic assessment of AMR a) Longitudinal metaproteomic assessment of AMR within the WTTP. b) metagenomic and metatranscriptomic normalized protein levels linked to AMR within the WTTP over time. c)Tripartite network assessing the normalized protein levels derived from MGEs and associated taxa. Boxplots depicting significant differential (*adj.p < 0.05,* Two-way ANOVA) abundance of aminoglycoside resistance in plasmid versus phage in *Candidatus* Microthrix parvicella as well as overall.

## Discussion

The surveillance of wastewaters for the identification of microbial molecular factors is a critical tool for identifying potential pathogens. This has been highlighted recently with the tracking of SARS-CoV-2 within wastewater treatment plants to assess viral prevalence and load within a given community^35^. Such approaches have also been employed for screening for antimicrobial resistance at a population level^36,37^. So far, several studies^10,16,38,39^ have characterized the proliferation of ARGs and antibiotic resistant bacteria in BWWTP. Szczepanowski *et al.*^38^ identified 140 clinically relevant plasmid-derived ARGs in a BWWTP metagenome^38^ while Parsley *et al.*^39^ characterized ARGs from bacterial chromosomes, plasmids and in viral metagenomes found in a BWWTP^39^. Further studies have shown that conventional BWWTP processes at best only partially remove ARGs from the effluent and may find their way into the urban water cycle^40–42^. Wastewater treatment plants, therefore, are crucial reservoirs of AMR, whose monitoring may allow for early-detection of AMR within the human population feeding into the system. Here, we leveraged a systematic and longitudinal sampling scheme from a BWWTP to identify diverse AMR categories prevalent within the BWWTP microbial community. In line with the studies by Szczepanowski *et al.*^38^ and Parsley *et al.*^39^, we found up to 29 AMR categories with several ARGs within the BWWTP. More importantly, and unlike the previous studies, we linked the identified ARGs to clinically-relevant ESKAPEE pathogens, which represent a growing global threat to human health.

In our BWWTP samples, we identified a core group of 15 AMR categories that were ubiquitous at all timepoints. In line with the above-mentioned reports, the observed core resistance categories may reflect their abundance in the surrounding human population^43^. This has previously been reported by Pärnänen *et al.*^44^, Su *et al.*^45^ and Hendriksen *et al.*^16^ where they showed that BWWTP AMR profiles correlate with clinical antibiotic usage as well as other socio-economic and environmental factors. On the other hand, bacteria are known to have innate defense mechanisms against inhibitory bacteriocins from other taxa^46^. Therefore, one must be cognizant of the phenomenon that the observed core group of AMR categories may also be a proxy for the abundance of specific resistant bacteria. Despite this observation, it is plausible that both anthropogenic and microbial sources for AMR play a role in the observed resistance categories within the BWWTP. Interestingly, we found that several AMR categories, including ancillary (prevalent, moderate, and rare) groups, were associated with *M. parvicella* within the BWWTP. Similar to the findings by Munck *et al.*^47^, we found a wide range of bacteria associated with AMR categories including Acidimicrobiales, Burkholderiales and Rhodocyclales. On the other hand, we report that taxa, including ESKAPEE pathogens, belonging to 25 bacterial orders were associated with 29 AMR categories, compared to the eight bacterial orders reported previously.

It is important to note that the mobilome plays a critical role in the dissemination of AMR within microbial communities. AMR from resistant bacteria within the BWWTP can quickly disseminate within the BWWTP^11,25^, including transmission from pathogenic to commensal species^48,49^. As a result, mediated through HGT, the BWWTP becomes a hotspot for resistant bacteria, which are then released back into the receiving environment^50^, and eventually the human population^11,51^. Therefore, to limit the dissemination of AMR, it is important to understand the role of MGEs within the BWWTP. Our comprehensive analyses identified the differential contributions of AMR transmission mediated via phage and plasmid (Fig. 7). Specifically, we identified clear segregation of aminoglycoside, bacitracin, MLS and sulfonamide resistance categories with plasmids, while fosfomycin and peptide resistance were increasingly encoded and conferred via phages. While the association between these AMR categories and plasmids^52–55^ or phages^56^ are in line with previously reported results, differential analysis between MGEs has not been previously reported and has not been performed on multi-omic levels. As such, in this study we report for the first time the systematic and extensive comparison of AMR encoded and expressed by phages versus plasmids. Our results indicating the segregation of ARGs within the ESKAPEE taxa via the MGEs further provide insights into potential modes of AMR transmission among pathogens. Though one cannot exclude the possibility of transmission of the above-mentioned ARGs via other MGEs, identifying potential segregation of MGEs in the transmission of ARGs brings us one step closer to identifying specific transmission paths and limiting the spread of AMR. For example, some studies have reported plasmid “curing”, the process by which plasmids are removed from bacterial populations, as a strategy against dissemination of AMR^57,58^. As described by Buckner *et al.*^59^ plasmid curing, as well as other anti-plasmid strategies, could both reduce AMR prevalence, and (re-)sensitize bacteria to antibiotics^59^. Combining these strategies with AMR categorization according to preference for specific MGEs will give us novel strategies for removing MGE-mediated resistance in the fight against AMR.

**Figure 7.**
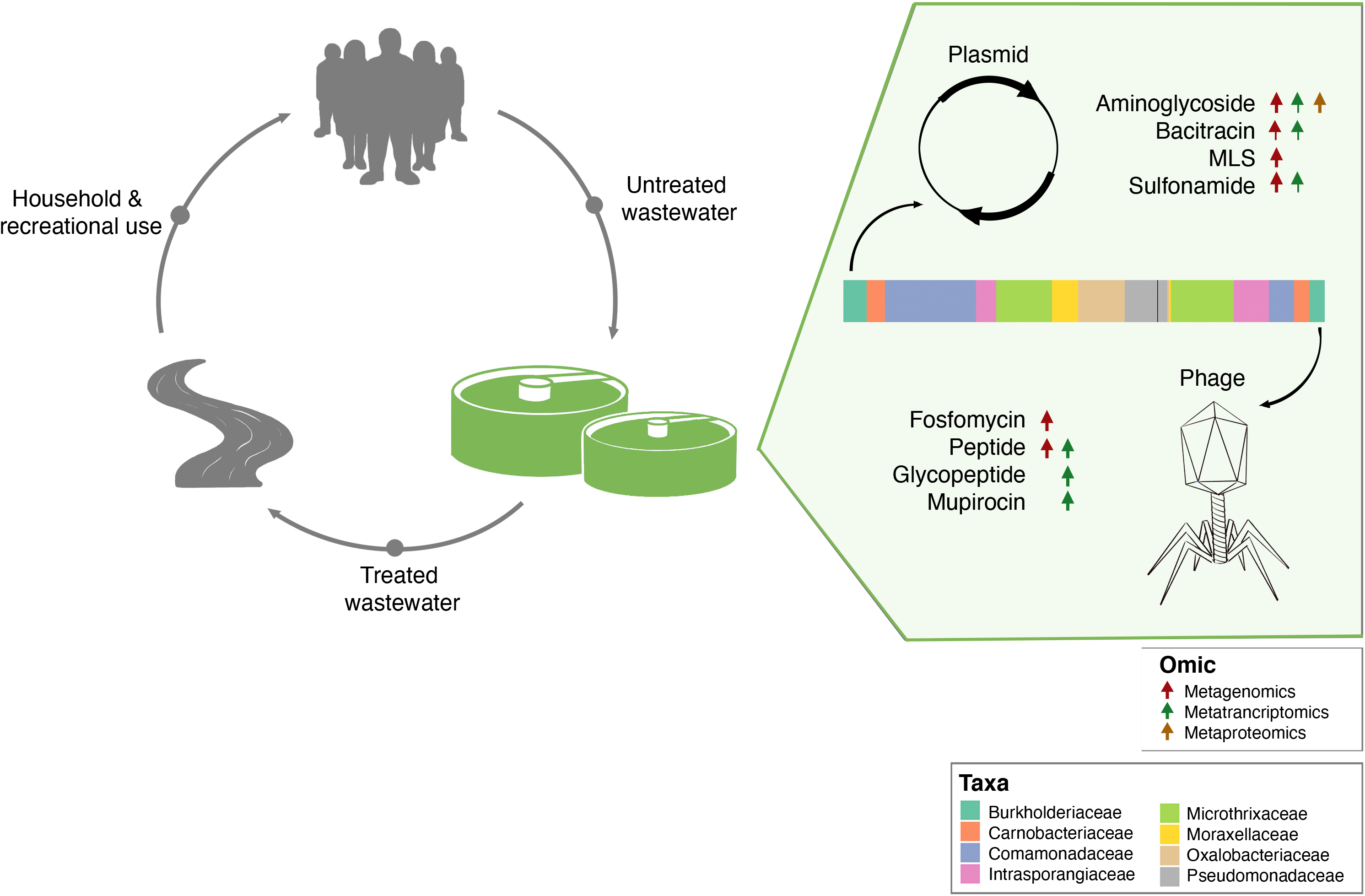
Separation of MGE-derived AMR within the BWWTP. A graphical summary highlighting AMR categories found significantly increased in phage versus plasmid in all three omes.

By complementing the metagenomic analyses, metatranscriptomics conferred essential information regarding gene expression within the resistome. For instance, when comparing AMR expression levels of aminoglycoside, bacitracin, and sulfonamide mediated via MGEs, it is noticeable that expression levels in plasmids mirror the genomic content, i.e. they exhibited higher levels of expression when compared to phage. On the other hand, glycopeptide and mupirocin resistance genes which were highly expressed in phages were not reflected within the metagenomic data. Additionally, we found the *YojI* resistance gene to be more highly expressed than any other ARGs. To facilitate resistance against the peptide antibiotic microcin J25, the outer membrane protein, TolC, in combination with *YojI* is required to export the antibiotic out of the cell^29^. Microcin J25 belongs to the group of ribosomally synthesized and post-translationally modified peptides (RiPPs) and has antimicrobial activity against pathogenic genera such as *Salmonella* spp. and *Shigella* spp.^60^. Interestingly, it has only recently been proposed as a treatment option against *Salmonella enterica and* has been discussed in recent years as a potential novel antibiotic^61^. Based on these results, by considering that BWWTPs may reflect both the presence of AMR within the human population as well as be a hotspot of dissemination and generation of new AMR, surveillance of BWWTPs must be emphasized when developing new antibiotics. Our findings collectively suggest that the differential capacity of MGEs to disseminate AMR, coupled with longitudinal and expression-level analyses are crucial for monitoring human health conditions. More importantly, we report for the first time that BWWTP monitoring for AMR may allow for early detection of previously undescribed and previously undescribed resistance mechanisms.

Finally, we applied an integrated multi-omic approach to improve our knowledge on the functional potential of AMR and simultaneously validate the abundance and expression findings of the ARGs. By normalizing the metaproteomic results with the normalized expression of genes we were able to assess the stability of expressed AMR across time. We find that our methodology allows for an unbiased assessment of overall expression accounting for gene copy abundance and expression. These findings support the notion that the AMR genes may serve as sentinels or indicators of the presence of particular antimicrobial agents. However, it is plausible that we are only identifying the most abundant proteins and/or proteins that are more stable over time, and do not capture the entirety of the proteome profiles. Factors such as protein decay rates^62^ among others, may additionally influence this assessment. Irrespective of these observations, we identified segregation of AMR categories with respect to plasmids and phages. Our findings also highlighted the potential for identifying segregation of AMR via specific MGEs with an aim towards possible therapeutic and mitigation strategies via for example plasmid curing. Furthermore, we demonstrate that longitudinal analyses are required to survey AMR within BWWTPs due to the variations in the resistome across time. These shifts may either be representative of a shift within the human population itself, which in turn could be associated with the concurrent use of antibiotics at a given time, or competition within the microbial community. In any case, an independent or static analysis of the various time points may show an incomplete view of the BWWTP resistome, thus underlining the importance of our longitudinal resistome analyses. Overall, our findings suggest that BWWTPs are critical reservoirs of AMR, potentially allowing for early detection and monitoring of pathogens and novel resistance mechanisms linked to the introduction of new antimicrobials, whilst serving as a model for understanding the separation of MGEs through AMR.

## Methods

### Sampling and biomolecular extraction

From within the anoxic tank of the Schifflange municipal biological wastewater treatment plant (located in Esch-sur-Alzette, Luxembourg; 49° 30’48.29”N; 6° 1’4.53”E) individual floating sludge islets were sampled according to previous described protocols^28^. Sampling was performed starting on 21-03-21 till 03-05-2012 in approximately one-week intervals resulting in a total of 51 samples. DNA, RNA and proteins were extracted from the samples in a sequential co-isolation procedure as previously described^63^.

### Sequencing and data processing for metagenomics and metatranscriptomics

Paired-end libraries were generated for metagenomics with the AMPure XP/Size Select Buffer Protocol following a size selection step recommended by the standard protocol. Libraries for metatranscriptomics were prepared from RNA after washing stored extractions with ethanol and depletion of rRNAs with the Ribo-Zero Meta-Bacteria rRNA Removal Kit (Epicenter). Subsequently, the ScriptSeq v2 RNA-seq library preparation kit (Epicenter) was used for cDNA library preparation, followed by sequencing on an Illumina Genome Analyses IIx instrument with 100-bps paired-end protocol. Processing and assembly of metagenomic and metatranscriptomic reads was done using the Integrated Meta-omic Pipeline^64^ (IMP v1.3; available at https://r3lab.uni.lu/web/imp/). For the IMP processing, Illumina Truseq2 adapters were trimmed, and reads of human origin were filtered out, followed by a de novo assembly with MEGAHIT^65^ v1.0.6. Both metagenomic and metatranscriptomic reads were co-assembled to increase contiguity of the assemblies^64^.

### Identification of antimicrobial resistance genes and association with mobile genetic elements

The assembled contigs from IMP were used as input for PathoFact^66^, for the prediction of antimicrobial resistance genes, and to annotate MGEs. ARGs were further collapsed into their respective AMR categories, as identified by PathoFact in accordance with those provided by the Comprehensive Antibiotic Resistance Database (CARD)^67^. Thereafter, the raw read counts per ORF, as given by PathoFact, were determined with featureCounts. The relative abundance of the ARGs was calculated using the RNum_Gi method described by Hu *et al.*^68^ This method was applied to the BAM files generated by mapping using bwa^69^ which were further processed by samtools^70^, both for the metagenomic and metatranscriptomic reads independently, to extract gene copy number and transcriptome expression respectively, per sample.

Identified ARGs and their categories were further linked to associated bacterial taxonomies using the taxonomic classification system Kraken2^71^. Kraken2 was run on the contigs using the maxikraken2_1903_140GB (March 2019, 140GB) (https://lomanlab.github.io/mockcommunity/mc_databases.html)database71. Furthermore, utilizing PathoFact, AMR genes were linked to predicted mobile genetic elements (i.e. plasmids and phages) to track transmission of AMR between taxa. Specifically, to link both the MGEs and the taxonomy to the AMR genes, we mapped the genes to assembled contigs. By considering all different predictions of MGEs, a final classification was made based on the genomic contexts of the AMR genes encoded on plasmids, phages or chromosomes, including classification of those that could not be resolved (ambiguous). The AMR genes that could not be assigned to either the MGEs or bacterial chromosomes were subsequently referred to as *unclassified* genomic elements.

### Metaproteomics and data analyses

Raw mass spectrometry files were converted to MGF format using MSconvert^72^ with default parameters. The metaproteomic searches were performed using SearchGUI / PeptideShaker^73^ for each time point. To generate the databases, each predicted protein sequence file was concatenated with the cRAP database of contaminants (*common Repository of Adventitious Proteins*, v 2012.01.01; The Global Proteome Machine) and with the human UniProtKB Reference Proteome^74^. In addition, inversed sequences of all protein entries were concatenated to the databases for the estimation of false discovery rates (FDRs). The search was performed using SearchGUI-3.3.20^75^ with the X!Tandem^76^, MS-GF+^77^ and Comet^78^ search engines using the following parameters: Trypsin was used as the digestion enzyme and a maximum of two missed cleavage sites was allowed. The tolerance levels for identification were 10 ppm for MS1 and 15ppm for MS2. Carbamidomethylation of cysteine residues was set as a fixed modification and oxidation of methionines was allowed as variable modification. Peptides with a length between 7 and 60 amino acids and with a charge state composed between +2 and +4 were considered for identification. The results from SearchGUI were merged using PeptideShaker-1.16.45^73^ and all identifications were filtered in order to achieve a peptide and protein FDR of 1%.

Each predicted protein sequence corresponded to the predicted ORFs generated by the Prodigal (version 2.6.3) predictions included in PathoFact. As such predicted protein sequences matched the ARG annotation of the ORFs as provided by PathoFact.

### Multi-omic integration

To further improve upon the understanding of the AMR expression and assess its stability across time, we estimated the normalized protein index (NPI) per gene, by integrating the multi-omic data. To estimate the NPI, we first normalized the metaT abundance based on per gene copy numbers obtained via the metagenomic abundance:

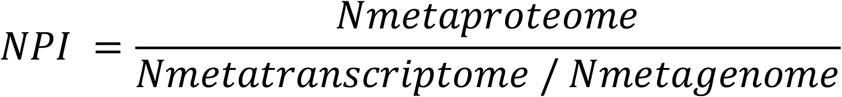

This, the normalized expression of genes, yields the per copy expression of ARGs within each AMR category. Subsequently, the normalized expression was used to standardize the metaP abundances for those genes where the necessary data was available.

### MGE partition assessment

To assess the segregation of MGEs through AMR we determined niche regions and overlap using the *nicheROVER* R package^79^. *nicheROVER* uses Bayesian methods to calculate niche regions and pairwise niche overlap using multidimensional niche indicator data (i.e. stable isotopes, environmental variables). As such, using AMR as the indicator data, we extended the application of *nicheRover* to calculate the probability for the size of the niche area of one MGE inside that of the other, and vice versa. We calculated the segregation size estimate for each MGE and additionally generated the posterior distributions of μ (population mean) for each AMR category in all omics. We further computed the niche overlap estimates between MGEs with a 95% confidence interval over 10 000 iterations.

### Data analysis

Figures for the study including visualizations derived from the taxonomic and functional analyses were created using version 3.6 of the R statistical software package^80^. A paired two-way ANOVA (Analysis of Variance) within the *nlme* package was used for identifying statistically significant differences for the AMR and taxonomic analyses. Tripartite and Bipartite networks were generated using the *SpiecEasi* ^81^ R package where a weighted adjacency matrix was generated using the Meinhausen and Buhlmann (*mb*) algorithm, with a nlambda of 40, and lambda minimum ratio at 0.001. The analyses were bootsrapped with n=999 to avoid overfitting, autocorrelations and false network associations. The network was further refined, selecting for positive edges, with a degree greater than the mean-degree of the initial network. The *igraph*^82^ package was used in R to render the graphics for the network. All code for visualization and analysis is available at: https://git-r3lab.uni.lu/laura.denies/lao_scripts.

## Data availability

The genomic FASTQ files from this work are publicly available at NCBI BioProject PRJNA230567. Metaproteomic data is publicly available at the PRIDE database under accession number PXD013655.

## Code availability

The open-source tools and algorithms used for the data analyses are reported in the Methods section, including relevant flags used for the various tools. Additionally, custom code for further analysis and generation of the figures can be found at: https://git-r3lab.uni.lu/laura.denies/lao_scripts

## Acknowledgements

We are thankful for the assistance of Audrey Frachet Bour, Lea Grandmougin, Janine Habier, Laura Lebrun (LCSB) for laboratory support. We acknowledge the valuable input from Rashi Halder at the LCSB Sequencing Platform with respect to library preparation. The computational analyses were performed at the HPC facilities at the University of Luxembourg (https://hpc.uni.lu)^83^.

## Figures

Supplementary figure 1. Expression levels of individual ARGs Expression levels of individual ARGs over time within the BWWTP.

Supplementary figure 2. Taxonomic diversity of AMR The plot indicates the number of taxa (order level) in which the corresponding AMR categories are identified.

Supplementary figure 3. Partitioning of MGEs through AMR The boxplots indicate the niche sizes (left) for the MGEs (plasmids and phages) based on metagenomic assessment. Niche plots (right) reveal that plasmids tend to differentiate from phages based on their capacity to encode for aminoglycoside resistance.

Supplementary figure 4. Differential AMR abundance in MGEs The barplot reports the log2foldchange of AMR categories over time in MGEs (plasmid versus phage) in: a) the general microbial population, b) *M. parvicella,* c) *Pseudomonas* spp. And d) *Comamonas* spp.

Supplementary figure 5. Expression of AMR categories in MGEs The barplot reports the expression levels of AMR categories over time in MGEs (plasmid versus phage) in: a) Acidimicrobiales, and b) Burkholderiales.

Supplementary figure 6. AMR protein abundances Barplot depicting protein abundances of various AMR categories over time.

