## Supplementary Figures for "Mobilome-driven segregation of the resistome in biological wastewater treatment"

### Supplementary figure 1

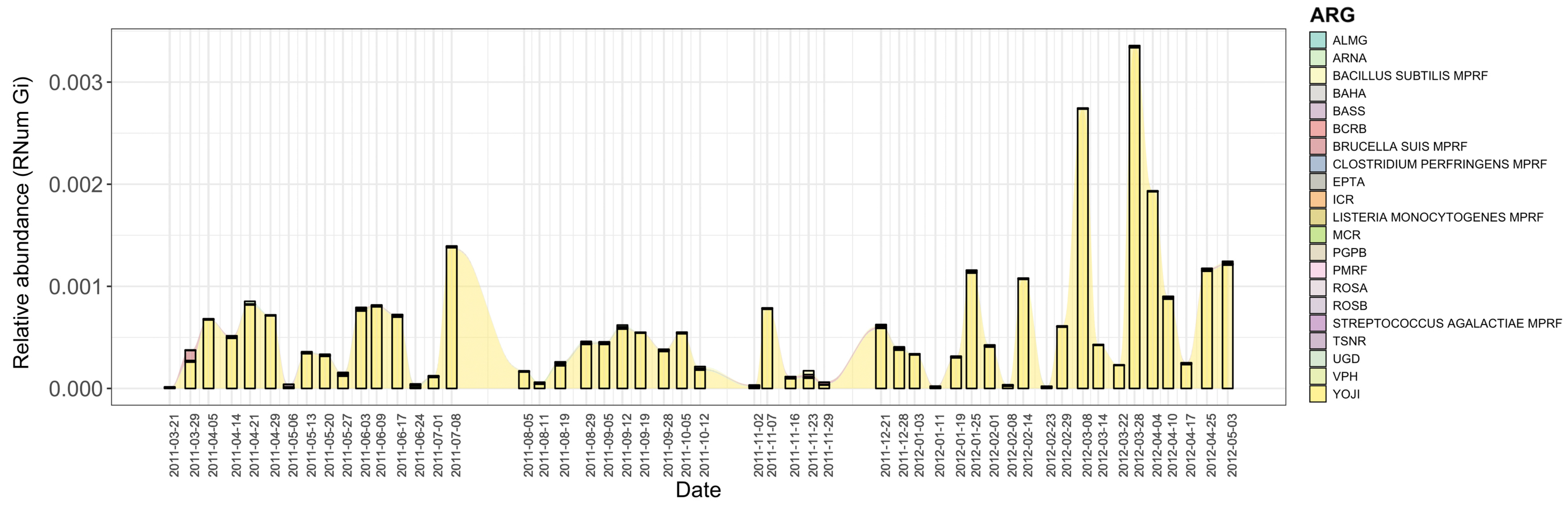


### Supplementary figure 2

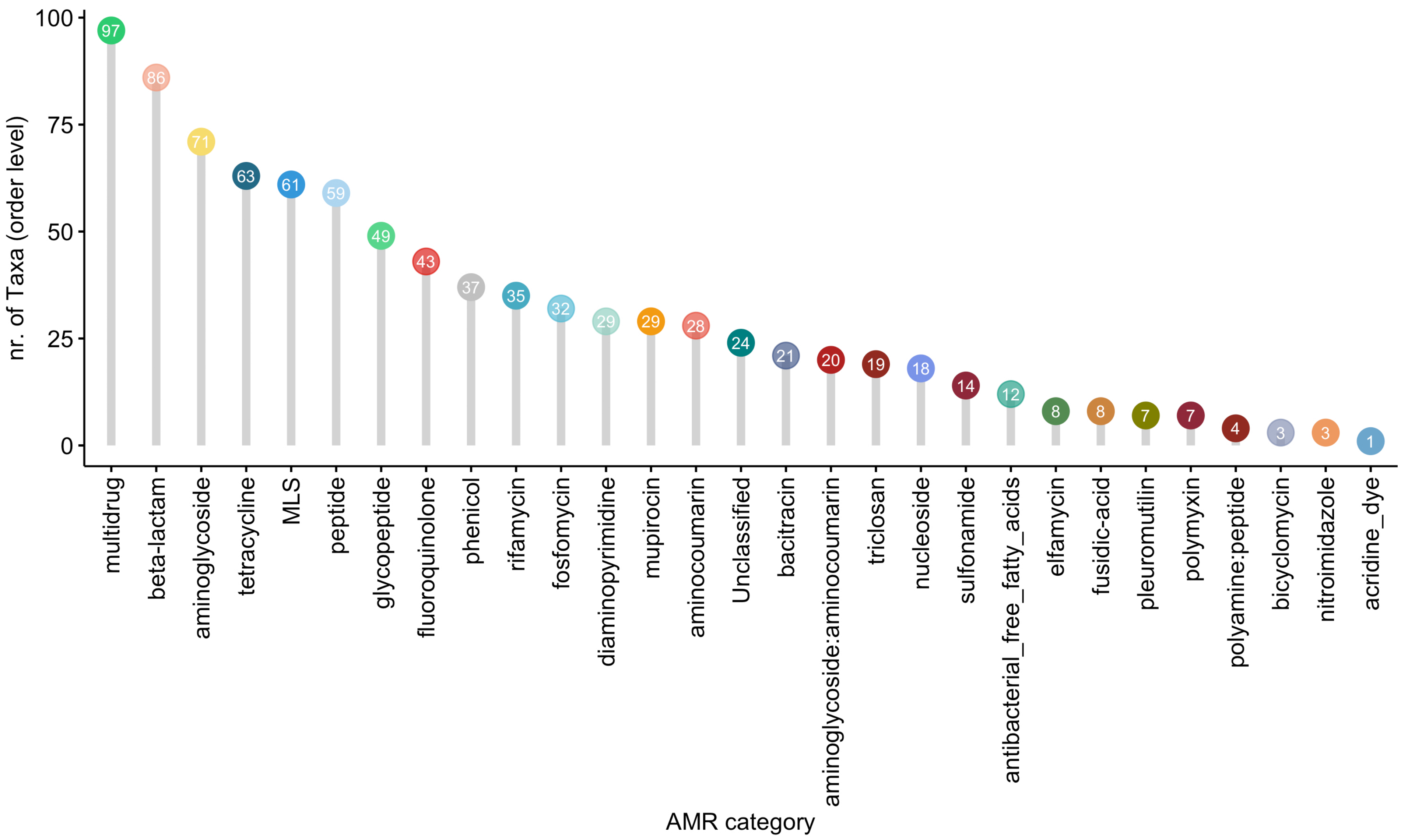


### Supplementary figure 3

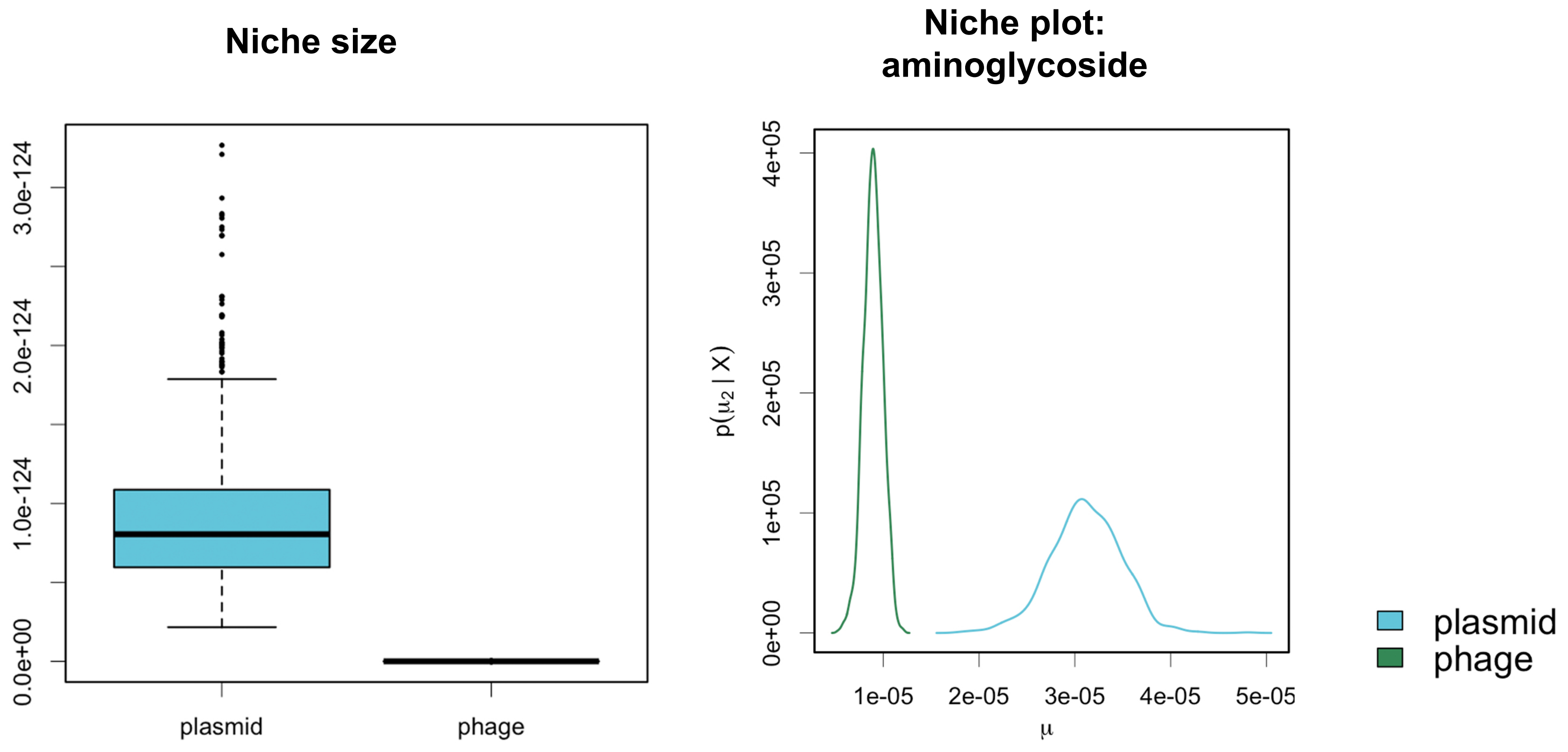

Supplementary figure 4

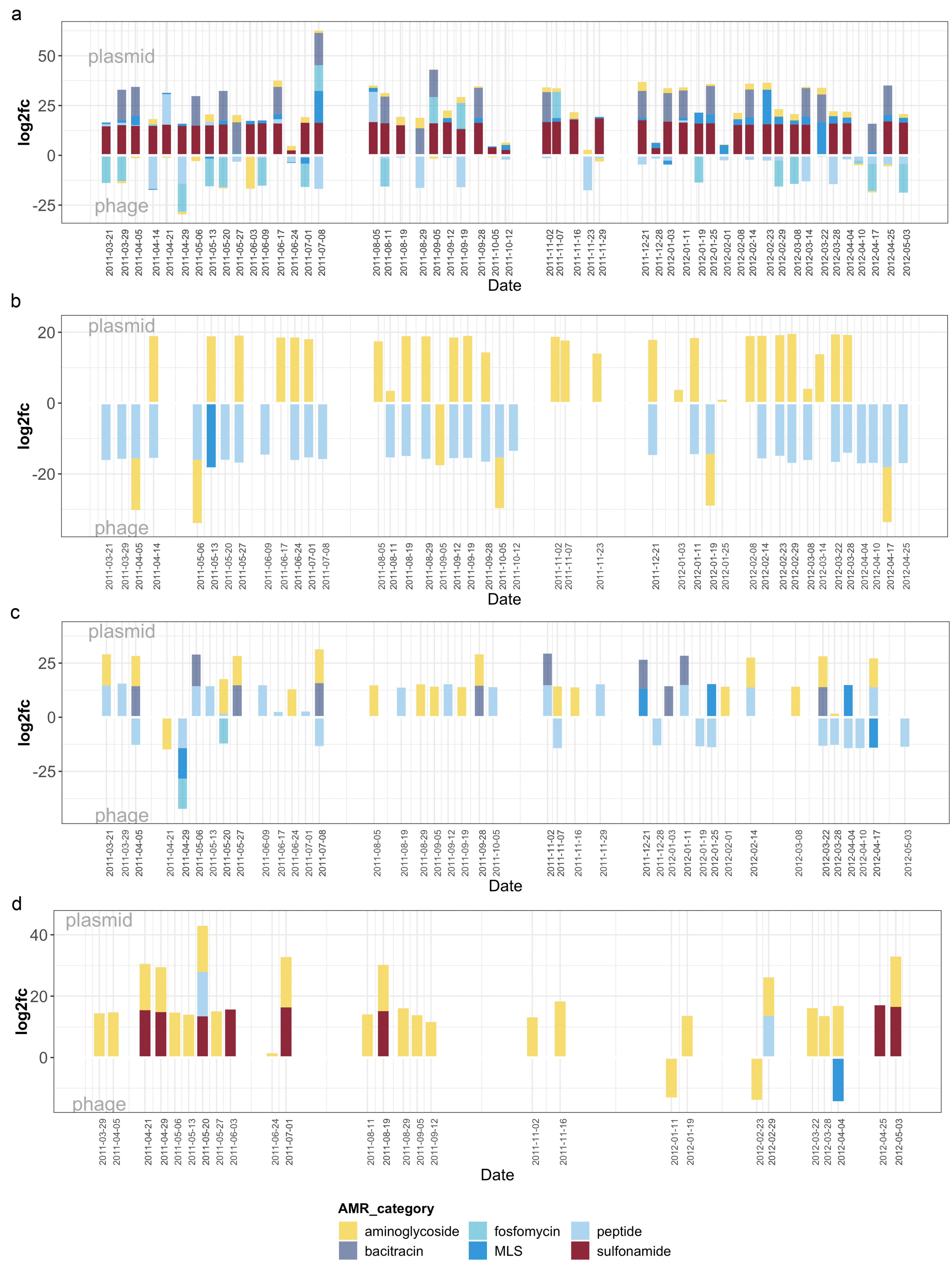

Supplementary figure 5

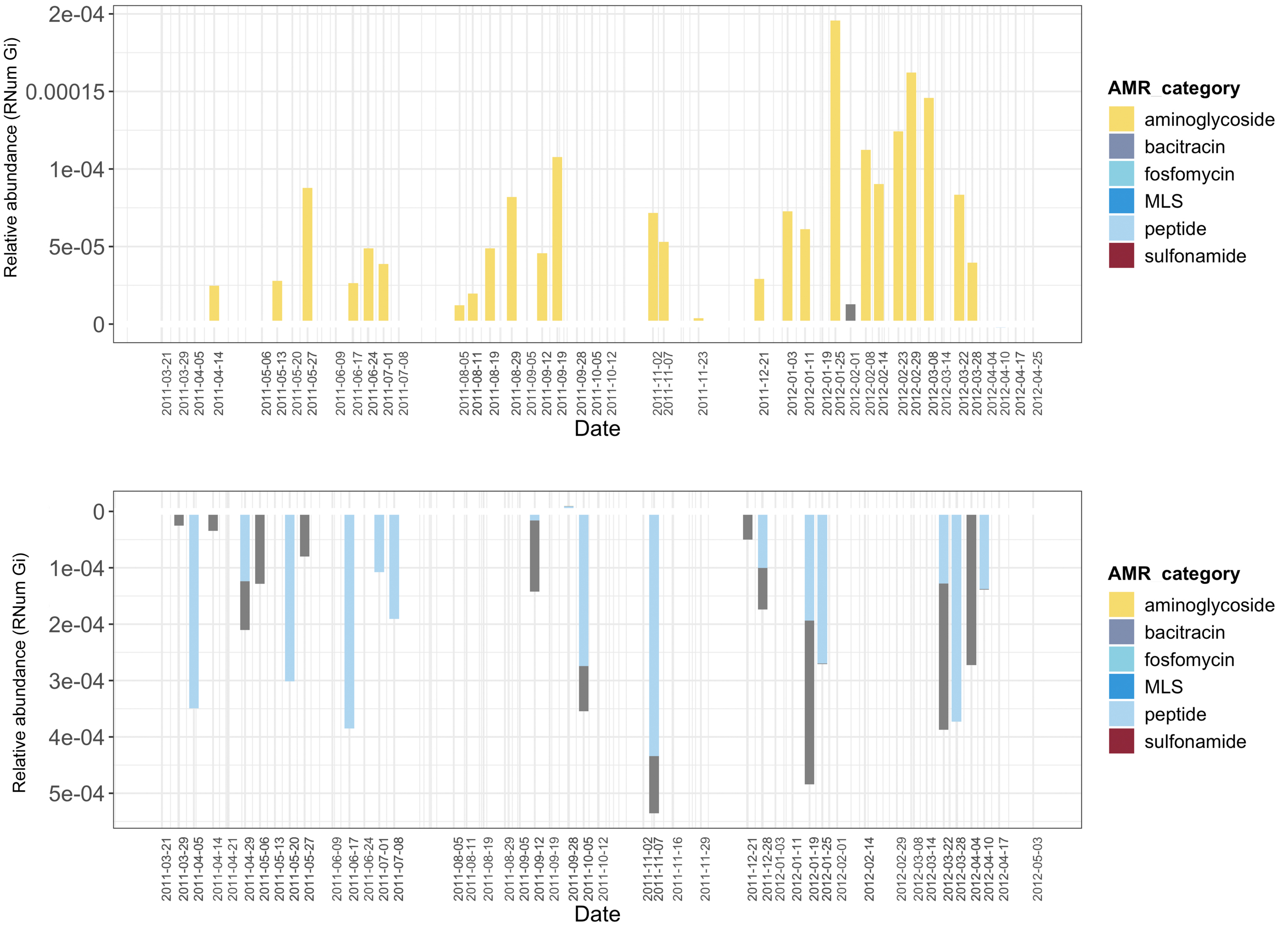


Supplementary figure 6

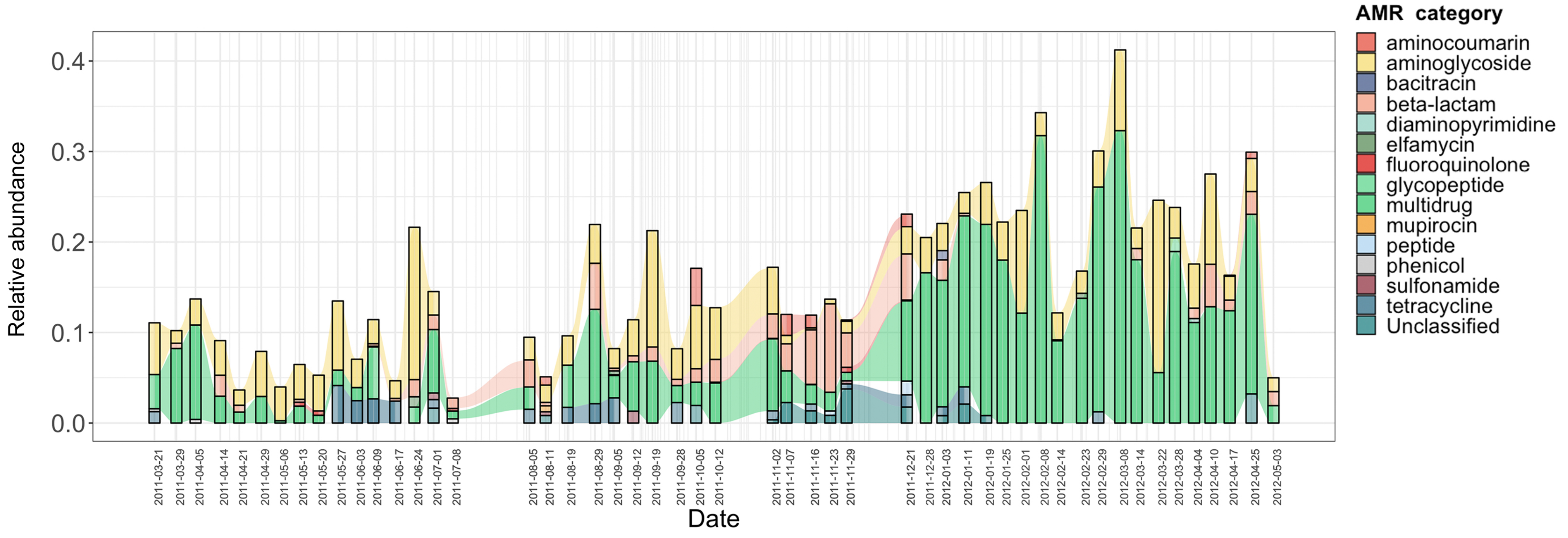
